# Towards a structurally resolved human protein interaction network

**DOI:** 10.1101/2021.11.08.467664

**Authors:** David F. Burke, Patrick Bryant, Inigo Barrio-Hernandez, Danish Memon, Gabriele Pozzati, Aditi Shenoy, Wensi Zhu, Alistair S Dunham, Pascal Albanese, Andrew Keller, Richard A. Scheltema, James E. Bruce, Alexander Leitner, Petras Kundrotas, Pedro Beltrao, Arne Elofsson

**Affiliations:** European Molecular Biology Laboratory, European Bioinformatics Institute (EMBL-EBI), Cambridge, UK; Science for Life Laboratory, Stockholm University 172 21 Solna, Sweden; Department of Biochemistry and Biophysics, Stockholm University, 106 91 Stockholm, Sweden; Biomolecular Mass Spectrometry and Proteomics, Bijvoet Center for Biomolecular Research and Utrecht Institute of Pharmaceutical Sciences, Utrecht University, 3584 Utrecht, The Netherlands; Netherlands Proteomics Center, 3584 Utrecht, The Netherlands; Department of Genome Sciences University of Washington Seattle WA 98109; Department of Biology, Institute of Molecular Systems Biology, ETH Zurich, Otto-Stern-Weg 3, 8093 Zurich, Switzerland; Center for Computational Biology, The University of Kansas, Lawrence, KS 66047, USA

**Author notes:** Contributed equally (David F. Burke, Patrick Bryant, Inigo Barrio-Hernandez, Danish Memon, Gabriele Pozzati).

## Abstract

All cellular functions are governed by complex molecular machines that assemble through protein-protein interactions. Their atomic details are critical to the study of their molecular mechanisms but fewer than 5% of hundreds of thousands of human interactions have been structurally characterized. Here, we test the potential and limitations of recent progress in deep-learning methods using AlphaFold2 to predict structures for 65,484 human interactions. We show that higher confidence models are enriched in interactions supported by affinity or structure-based methods and can be orthogonally confirmed by spatial constraints defined by cross-link data. We identify 3,137 high confidence models, of which 1,371 have no homology to a known structure, from which we identify interface residues harbouring disease mutations, suggesting potential mechanisms for pathogenic variants. We find groups of interface phosphorylation sites that show patterns of co-regulation across conditions, suggestive of coordinated tuning of multiple interactions as signalling responses. Finally, we provide examples of how the predicted binary complexes can be used to build larger assemblies. Accurate prediction of protein complexes promises to greatly expand our understanding of the atomic details of human cell biology in health and disease.

## Introduction

Proteins are key cellular effectors determining most cellular processes. These rarely act in isolation, but instead, the coordination of the diversity of processes arises from the interaction among multiple proteins and other biomolecules. The characterization of protein-protein interactions is crucial for understanding which groups of proteins form functional units and underlies the study of the biology of the cell. Diverse experimental and computational approaches have been developed to determine the protein-protein interaction network of the cell (i.e. the interactome) with hundreds of thousands of human protein interactions determined to date ^1–3^. These interactions vary from transient interactions that can regulate an enzyme to permanent interactions in large molecular machines.

The structural characterisation of any interactome has lagged behind its experimental determination due to technical limitations, with experimental and homology models currently covering an estimated 15 thousand human interactions ^4,5^. The structural characterisation of protein complexes is a critical step in understanding the mechanisms of protein function, studying the impact of natural and disease mutations ^4,6–8^ and the regulation of cellular processes via the post-translational tuning of binding affinities ^9–12^.

While there has been great progress in experimental techniques for determining large complexes, current experimental approaches are not easily scalable. Computational approaches for predicting the structure of interacting protein pairs on a large scale have relied primarily on identifying structural similarity for pairs of proteins against experimentally determined protein complexes ^4,6,13,14^. Based on these approaches, the Interactome3D^4,14^ repository currently lists 7625 predicted models based on homology of domains, a number similar to the 8359 interactions listed in this resource as having an experimentally determined model. In addition to modelling based on homology, co-evolution based information has been used to predict interaction pairs and guide structural docking for bacterial proteins ^15^. More recently, neural network-based approaches have demonstrated the ability not only to accurately predict the structures of individual proteins ^16,17^ but also the structure of protein complexes ^16,18–21^. In benchmark sets where protein pairs are known to form a direct complex, these approaches can correctly predict the structure of up to 60% of the dimers ^18^. These methods have been recently used to predict structures of 1,506 *S. cerevisiae* interactions that were selected based on co-evolution signals ^22^. However, the application of these neural network models for the large-scale prediction of human complex structures has not been tested yet.

Here, we assess the possibilities and limitations of applying AlphaFold2 to modelling human interactions on a large scale. We predicted the complex structures for two sets of human interactions obtained using different experimental methods, comprising 65,484 unique human interactions. We show that metrics derived from the predicted structures can be used to rank the models according to confidence, with 3,137 predicted structures ranked as highly confident. Further, we show that the higher confidence predictions are enriched among those supported by a combination of methods indicating or by constraints indicated by orthogonal cross-link data. We showcase the value of a structurally resolved interactome by studying disease mutations and phosphorylation of interface residues. Finally, we provide some indication that binary complexes can be used to build higher-order assemblies.

## Results

### Structure prediction of high confidence human protein interactions

We selected experimentally determined human interactions from the Human Reference Interactome (HuRI) and the Human Protein Complex Map (hu.MAP 2.0). HuRI comprises interactions determined by yeast two-hybrid screening ^2^ from which we modelled 55586 pairs. From hu.MAP we selected 10,207 high-quality (confidence score ≥0.5) protein-protein interactions (PPIs), which were derived by integration of affinity purification, co-fractionation and proximity ligation assays ^3^. While HuRI is more likely to be enriched for direct protein interactions, including potentially transient partners, the hu.MAP set is more likely to reflect stable protein interactions, including members of the same complex that may not necessarily interact directly. The overlap between the two datasets is small (309 pairs), and a comparison with two large scale compendiums of structural models^4^, (see **Methods**) indicates that 62,019 of the combined pairs do not have experimental models or can be modelled easily by homology, suggesting a significant potential gain in structural knowledge.

We predicted the structure of 65,484 non-redundant pairs using the FoldDock pipeline ^18^, based on AlphaFold2 ^17^. We have previously shown that larger interface size and higher predicted lDDT (plDDT) scores from AlphaFold2 in the interfaces of the predicted complexes are associated with more reliable predictions ^18^. As in the FoldDock pipeline, we combined these two metrics into a single score, which can be used to predict the DockQ score of a complex, dubbed pDockQ (**Methods**) that can rank models by confidence. We tested the overlap performance and ranking by pDockQ score by comparing the predicted models with experimental models. Across 1,465 comparisons, 742 (50%) of predicted complexes were deemed to be well modelled (DockQ>0.23). For predictions with pDockQ>0.23, we estimate that 70% (671 out of 955) are well modelled and for pDockQ>0.5, 80% (521 out of 651) are considered well modelled. However, it is worth noting that this estimated performance applies to cases where we expect that two proteins interact via a direct contact.

We show in **Fig. 1A** the distribution of pDockQ for the predicted interactions and a set of predicted structures for 2,000 random pairs of proteins. The pDockQ of known interacting proteins tends to be higher than for the random set with the predictions for hu.MAP showing on average higher confidence than the HuRI set. Additionally, when selecting hu.MAP interactions also supported by yeast-two-hybrid (Y2H) or cross-link data (cross-linking) results in much higher confidence values (**Fig. 1A**). This suggests that high confidence models are enriched for protein interactions supported by the two types of methods associated with high affinity and direct interactions. Based on the benchmarking results, we selected 3,137 structures (**Fig. 1B**) as high confidence models based on a cutoff of pDockQ>0.5, which would indicate around 80% of correct models compared to the experimental models. The number of structures increased to 10,061 if a cutoff of 0.23 is used. Only 0.3% of the random set of models would be considered a confident prediction at this cut-off. In **Fig. 1C** we show examples of predicted structures aligned to experimental or homology models, showing how the predictions and the confidence score relate to the observed alignments. For the majority of these cases, even with lower confidence values, the interaction interface is generally in good agreement, except for the interaction between subunits of the proteasome 26S complex, ATPpase domain 2(PSMC2) and non-ATPase domain 11 (PSMD11), which has the lowest confidence score of the illustrated models. It can be noted that several of the models in **Fig 1C** are parts of large complexes; PRDX2-PRDX3: members of the peroxiredoxin family of antioxidant enzymes, RFC2-RFC5: subunits of heteropentameric Replication factor C (RF-C), YWHAB-YWHAG: parts of the 14-3-3 family of proteins tyrosine 3-monooxygenase/tryptophan 5-monooxygenase activation proteins beta (YWHAB) and gamma (YWHAG), and RPL9-RPL18A: ribosomal proteins L99RPL9) and L18a (RPL18A). This shows that AlphaFold2 can predict the structure of directly interacting protein pairs present in large complexes.

**Figure 1.**
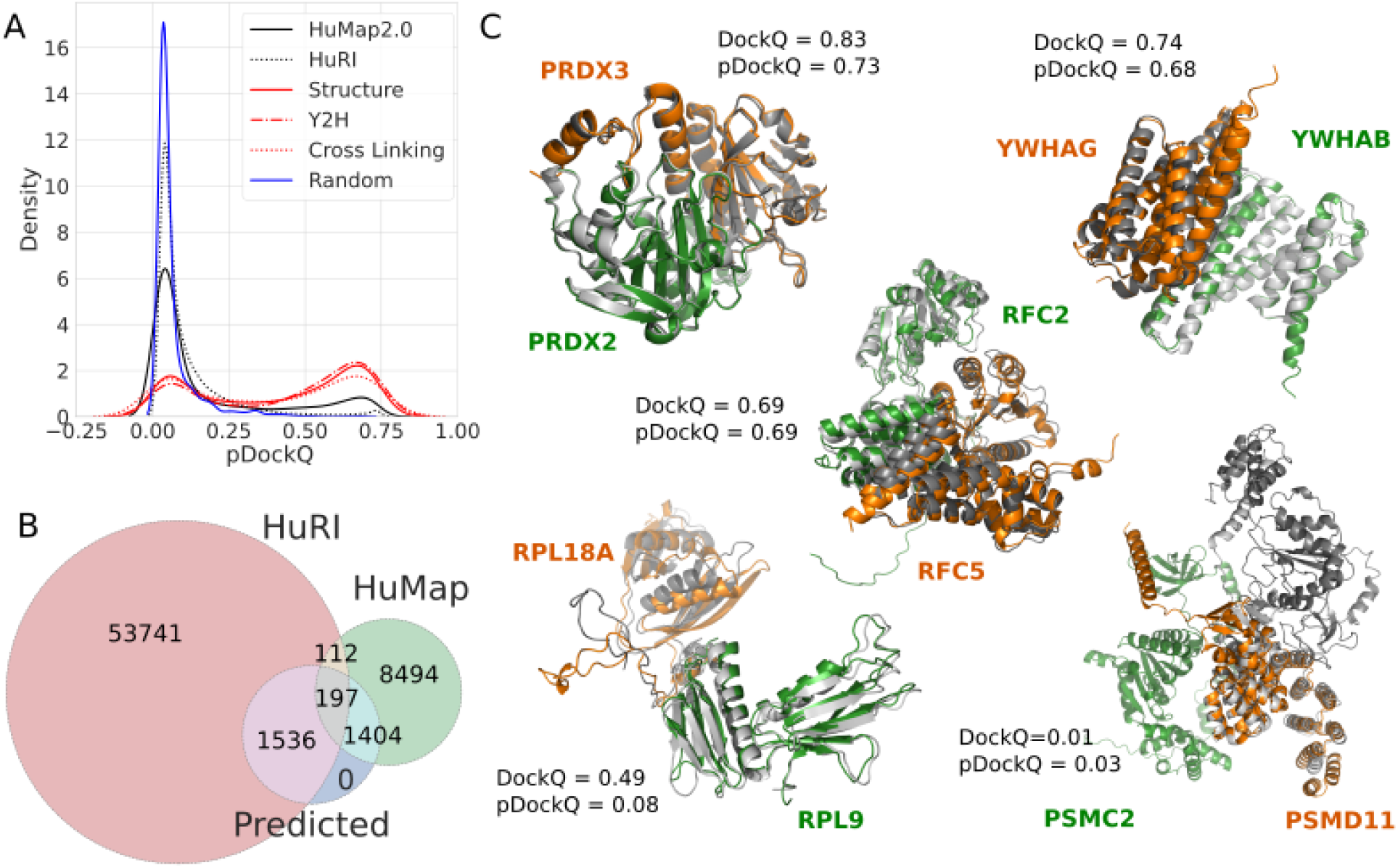
Application of AlphaFold2 complex predictions to a large dataset of human protein-protein interactions. A) Distribution of model confidence score (pDockQ) for predicted structures from two large human protein interaction datasets (hu.MAP and HuRI), compared with confidence metrics from 2000 random pairs of proteins. The hu.MAP dataset was further subsetted to those that have support from yeast two-hybrid (Y2H), cross-link data (Cross-linking) or correspond to pairs with available experimental or homology modelling information (Structure). B) Number interactions with models built from both datasets and those that we consider to be of high confidence (Predicted), corresponding to those with pDockQ>0.5. C) Examples of predicted models (orange and green) overlapped with the corresponding experimental models (grey) and the observed (DockQ) or predicted (pDockQ) quality of the models.

A list of protein interactions with predicted structural models, confidence metrics and annotations is provided in **Table S1**, and all models are available as described in the data availability section.

### Protein and interaction features impacting on prediction confidence

As shown in **Fig. 1A**, protein pairs present in PDB are enriched in high-scoring models compared with all other pairs in HuRI and Hu.MAP. There could exist several possible explanations for this, such as the inability of AlphaFold2 to identify transient or indirect interactions that might be more present in HuRI and Hu.MAP. Nevertheless, it is also possible that the two high-throughput datasets contain a fraction of non-interacting pairs. Therefore, to understand this difference better, we studied an additional dataset created from large (>10 chains) heteromeric protein complexes.

The set of large complexes consists of 12 large heteromeric protein complexes, and all (non-identical) pairs of protein chains in each complex were docked with each other. These pairs can be divided into the ones with direct interaction and those that do not. Here, we used a definition of more than 20 contacts less than 8Å between Calphas to exclude small interaction interfaces. When a complex contains multiple copies of identical chains all interactions were included to allow for alternative interactions between the chains. The difference in pDockQ scores between the direct and indirect interacting pairs is striking, where only 6% of the indirect pairs have a pDockQ score >0.5 compared with 38% of the directly interacting pairs (**Fig. 2A)**, showing that directly interacting pairs often can be predicted even when they are part of large complexes. However, as expected, AlphaFold2 models of members of the same complex that do not interact are assigned a low confidence.

**Figure 2:**
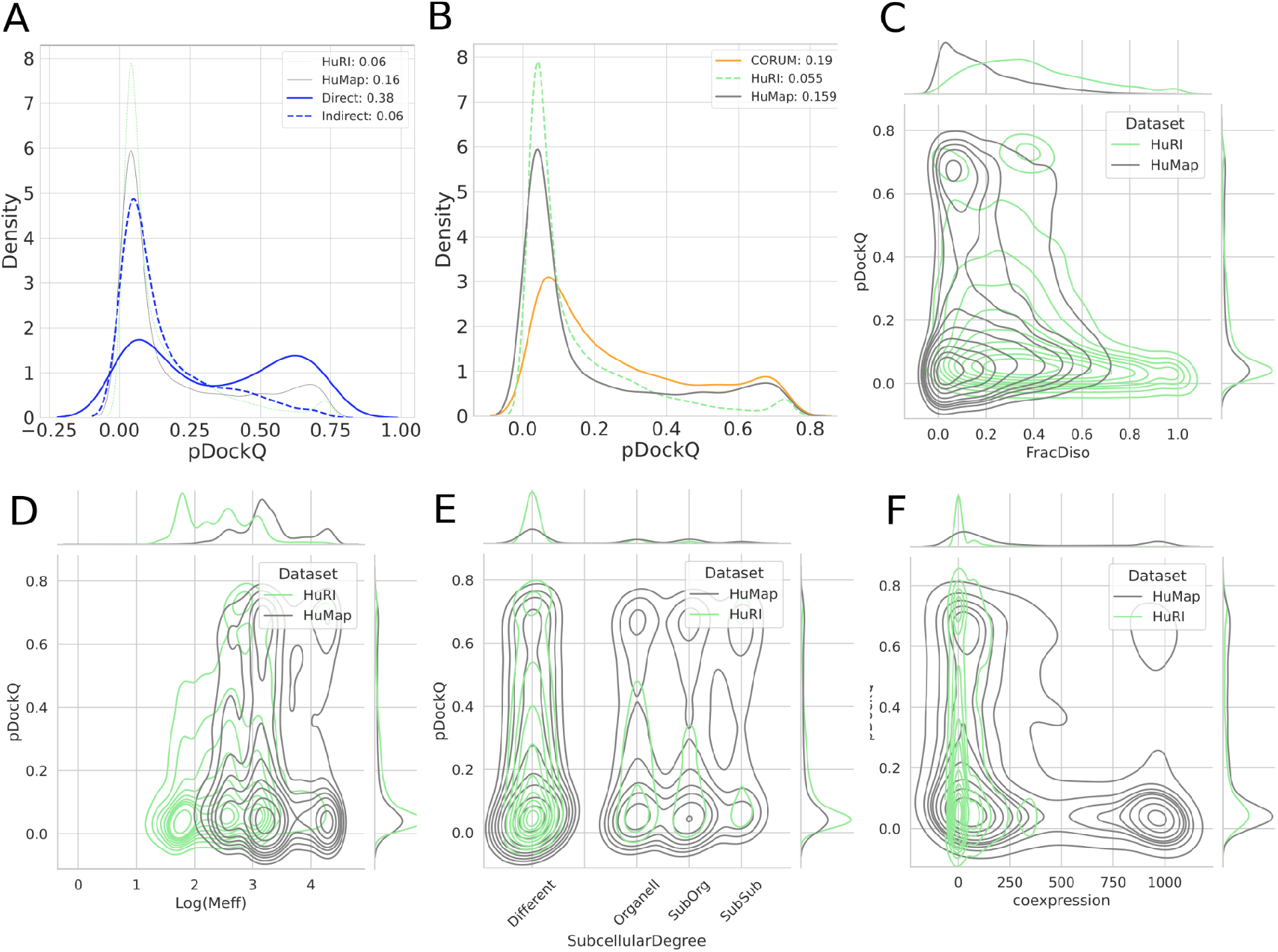
Protein and interaction features impacting on prediction confidence, analysis of different datasets. In all subfigures, proteins in HuRI in green, Hu.MAP grey, CORUM orange, and from large PDB complexes blue. A) pDockQ values of directly and indirectly interacting proteins from the same complex (blue), for comparison HuRI and Hu.Map data are shown with thin lines. B) pDockQ values of CORUM (orange), HuRI (green) and Hu.Map (grey) datasets. C) Fraction of residues predicted to be disordered (pLDDT<0.5) shows that protein pairs in HuRI are enriched in disorder. D) Proteins in HuRI have fewer sequences in the paired MSAs. E) Proteins that share subcellular localisation (solid lines) are enriched in high pDockQ scores in all three datasets. F) Only protein pairs in Hu.Map are coexpressed according to STRING and coexpressed pairs are enriched in pairs with high pDockQ scores.

In this study, we see that Hu.MAP has many more high-confident predictions than predictions from HuRI, which is based on yeast two-hybrid (Y2H) experiments. To further understand this difference, we first analyzed a subset of all protein pairs from the CORUM^23^ database, the best manually curated database of mammalian protein complexes. We selected a subset of the complexes and predicted the interaction of all pairs in the same complex. The average pDockQ score of CORUM is slightly higher than for Hu.MAP, but the number of high-quality predictions is similar (16% vs 19%), indicating that the different databases of protein complexes have a similar fraction of high-confidence predictions and that HuRI is an outlier **(Fig 2B)**.

It is unlikely that the Y2H in HuRI data should contain a large set of indirect interactions, as only two human proteins are expressed in the same cell. Therefore, there must be another reason for the few high-confidence predictions. We examined the properties of the pairs present in the two datasets. Here, it can be seen that HuRI proteins differ from the Hu.Map (and other datasets) in two ways. HuRI protein pairs contain more intrinsic disorder (**Fig 2C)** and have fewer sequences in their MSAS **(Fig 2D)**. In these figures it can also be seen that the pDockQ values tends to increase with less disorder and more sequences in the alignments, although it is clearly not an exact relationship. Further, protein pairs in HuRI are less likely to be found in the same subcellular compartment (**Fig 2E**), and have similar coexpression profiles **(Fig 2F**). Considering all this, it is likely that many interactions in HuRI are transient (or weak) and that AlphaFold2 cannot reliably predict such interactions. These factors also agree with our earlier study ^18^, where we showed that it is easier to predict interfaces containing secondary structure elements.

### Cross-linking support for predicted complex structures

Chemical cross-linking followed by mass spectrometry is an approach that can be used to identify reactive residues (usually lysines) that are in proximity, as constrained by the geometry of the cross-link agent used. The identification of such residues across a pair of proteins can help define the likely protein interface. To determine if the predicted complex structures agree with such orthogonal spatial constraints, we obtained a compilation of cross-links for pairs of residues across 528 protein pairs with predicted models (**Fig. 3A, Table S1**, see **Methods**). Of these, 51% of models have one or more cross-links at a distance below the expected maximal distance possible (**Fig. 3A, Methods**). Restricting the predicted models to higher confidence by the pDockQ score increases the fraction of complexes with cross-links within the maximal distance possible, reaching 75% for pDockQ scores greater than 0.5 (**Fig. 3A**). This result is in line with the benchmark results above, indicating that most models are likely to be correct at a pDockQ cut-off above 0.5. Additionally, predicted structures with pDockQ>0.23 are also likely to have many correct models as judged by the fraction supported by cross-linking.

**Figure 3.**
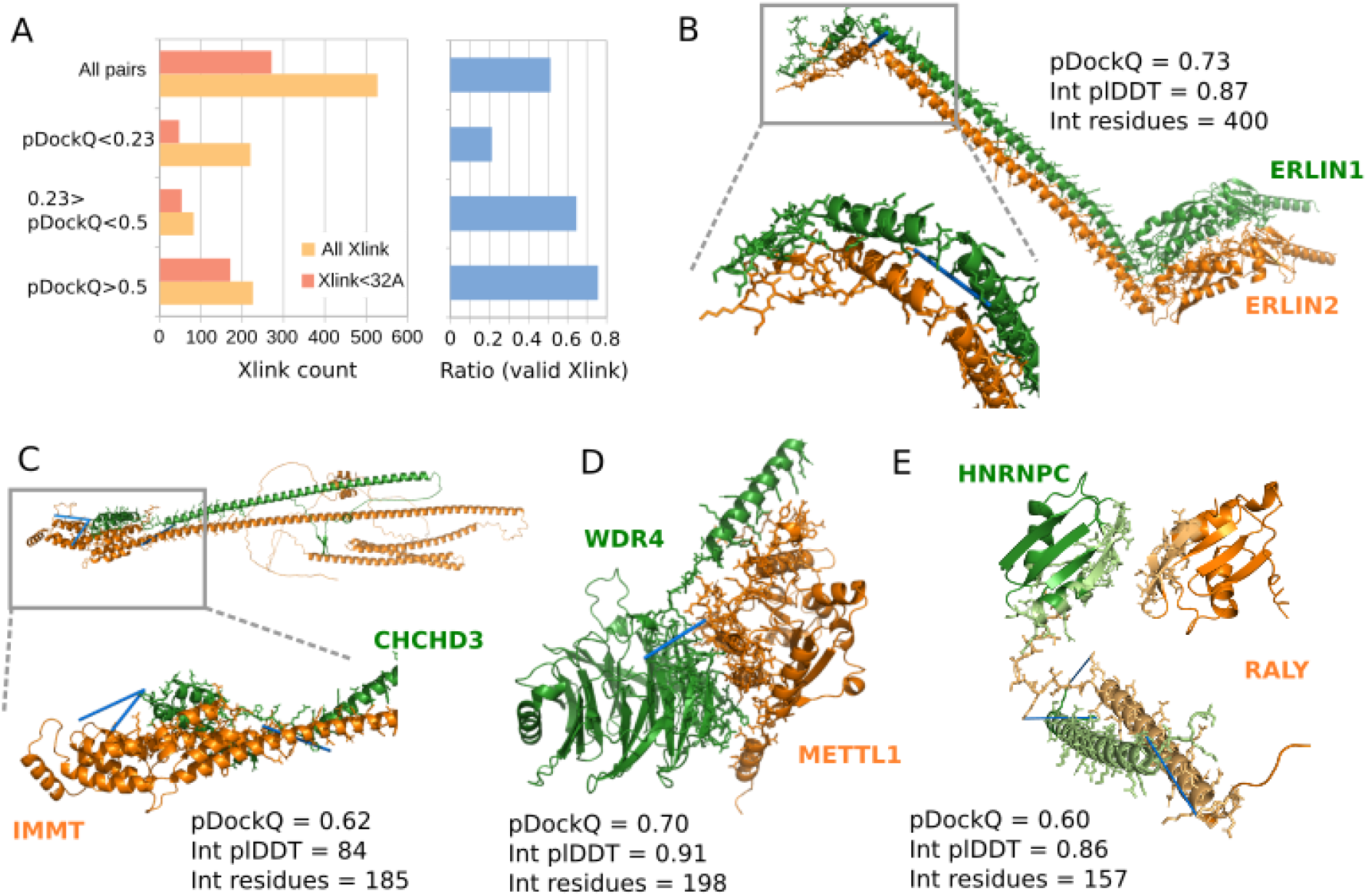
Cross-link support for predicted complex models. A) The numbers and ratios of predicted structures having cross-link information for pairs of residues that bridge the two proteins in the predicted structure, broken down by the cross-links that satisfy their expected maximal distance and by the predicted quality of the model (pDockQ). B-E) Examples of predicted structures of high confidence, no prior structural information, and at least supported by one cross-link (indicated with blue line).

In total, we have identified 479 cross-links providing supporting evidence for 171 predicted complex structures with pDockQ>0.5. Out of these, 41 correspond to complex structures with no experimental structure or homology models, from which we selected some to illustrate in **Fig. 3B-E. Fig. 23** shows the AF2 model for the full length of the ERLIN1/ERLIN2 complex, which mediates the endoplasmic reticulum-associated degradation (ERAD) of inositol 1,4,5-trisphosphate receptors (IP3Rs). AlphaFold2 predicts a globular domain (1-190) followed by an extended helical region with a kink around amino acid position 280. Unlike the model in Interactome3D, the paralogous proteins are stacked side-by-side with the hydrophobic face of the helices buried and the hydrophilic face (mainly Lys) exposed to solvent. A cross-link between the C-terminal residues K275 (ERLIN1) and K287 is predicted to bridge a distance of 18 Å, supporting the predicted model. In **Fig. 3C** we show the model for proteins IMMT and CHCHD3, components of the mitochondrial inner membrane MICOS complex. AlphaFold2 predicts a globular helical domain at the C-terminal end of IMMT (550-750) to interact with the C-terminal end of CHCHD3 (150-225). This is supported by data of three cross-links between; K173 (CHCD3) and K565 (IMMT), and K203 (CHCD3) to both K714 and K726 of IMMT. **Fig. 3D** shows the complex of tRNA-guanine-*N*(7)-methyltransferase (METTL) with its non-catalytic subunit (WDR4). The structure of WDR4 has not yet been solved experimentally but contains WD40 repeats, which are expected to form a β-propeller domain, as predicted here. The METTL domain is predicted to interact with the side of the WDR40, away from the ligand-binding pore. This orientation is supported by a cross-link between K122 (WDR4) and K143 (METTL) (18 Å).

Finally, in **Fig. 3E** we show the predicted complex structure for the heterogeneous nuclear ribonucleoprotein C (HNRNPC) and the RNA-binding protein, RALY. Two regions in both proteins are predicted with high confidence (plDDT>70), with the lower confidence regions not shown. The N-terminal domain in HNRNPC (16-85) is predicted to interact with the N-terminal domain of RALY (1-100). A long helix in HNRNPC (185-233) is predicted to interact with a helix in RALY (169-228). This interhelix interface is supported by cross-linking data for three pairs of lysines at either end of the helices (189→ 222; 229→179 and 232→ 183).

### Disease-associated missense mutations at interfaces

Missense mutations associated with human diseases can alter protein function via diverse mechanisms, including disrupting protein stability, allosterically modulating enzyme activity, and altering protein-protein interactions. Structural models can lead to the identification of interface residues allowing for the rationalisation of possible mechanisms of such interface disease mutations. To determine the usefulness of the predicted structures for studying disease mutations, we compiled a set of mutations located at interface residues that were previously experimentally tested for the impact on the corresponding interaction ^24^. We then performed *in silico* predictions of changes in binding affinity upon mutations using FoldX ^25^ and observed that mutations known to disrupt the interactions are predicted to have a strong destabilisation of binding compared to mutations known not to have an effect (**Fig. 4A, Table S2**). Very high confidence (plDDT>90) of the mutated residues led to more substantial discrimination between mutations known or not known to disrupt or not complex formation (**Fig. 4A**), indicating that only very accurate models are useful when using the FoldX forcefield for estimating the impact of binding affinity of mutations.

**Figure 4.**
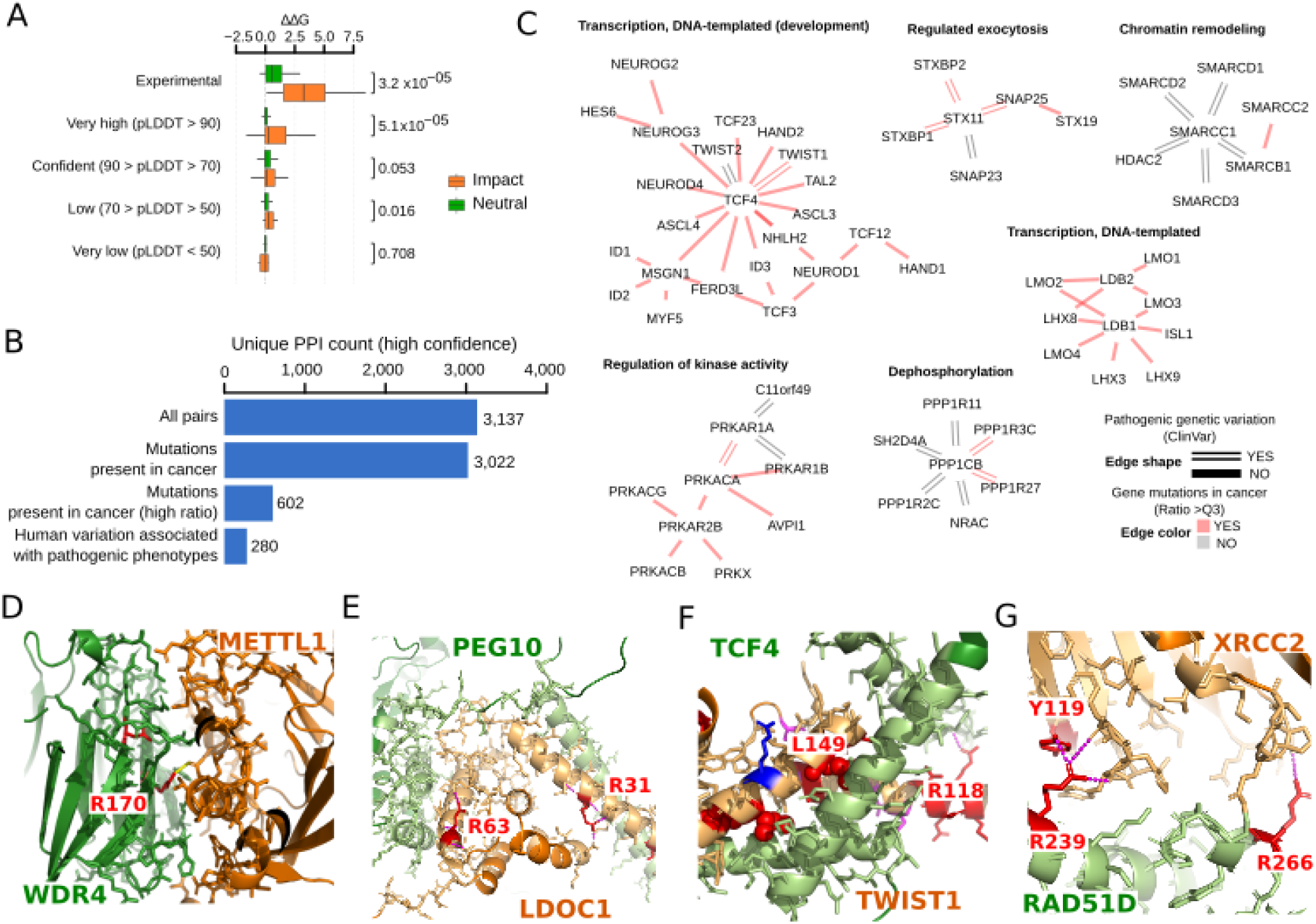
Disease mutations at protein complex interface residues. A) Boxplot showing the distribution of changes in predicted binding affinity (ΔΔG) for mutations known to have an impact (orange) versus the ones with neutral effect (green) B) unique protein-protein interaction pairs for high confidence models (pDockQ>0.5) in total, with mutations in cancer, mapped to the interface (all and top 25% ratios) and with pathogenic or likely pathogenic clinical variants mapped to the interface. C) Modules related to relevant biological processes. The colour of the edge represents the presence of cancer mutations in the interface (top25% ratio, colour red) and the shape of the presence of pathogenic clinical variants (double line). D-G) Selected relevant structures with no prior structural knowledge showing clinical variants or mutations in cancer mapped to the interface (mutated residues in red).

Having established the value of the predicted structures for modelling interface mutations, we mapped human disease (from ClinVar) and cancer mutations (from TCGA) to the interface residues defined by the set of high confidence protein complex predictions (pDockQ>0.5). The hu.MAP and HuRI confident predictions identified 280 interfaces carrying pathogenic mutations and 602 interfaces corresponding to the top 25% recurrently mutated interfaces in cancer, defined as the highest number of mutations per interface position (**Fig. 4B, Methods**). For these interface models we find a strong enrichment in pathogenic vs benign mutations at interface residues relative to the rest of the protein (2.3 fold enrichment, p-value 2.7×10^−31^)

We illustrate in **Fig. 4C** examples of protein network clusters with interface disease mutations across a range of biological functions. For example, interface mutations in chromatin remodelling, including members of SWI/SNF complex (SMARCD1, SMARCD2, SMARCD3) and several transcription factors related to the development (e.g. TCF3, TCF4, LMO1 and LMO2). All of the disease mutation information is provided in **Table S1**.

We selected examples of interfaces with disease mutations and no previous experimental data or homology to available models (**Fig. 4D-G**). **Fig. 4D** shows the interface of WDR4-METTL1 that has supporting cross-link information described above. WDR4 has two annotated pathogenic variants at this interface, linked with Galloway-Mowat Syndrome 6, with the highlighted R170 participating in interactions with a negatively charged residue of METTL1. **Fig. 4E** shows an example of an interface with 32 recorded interface mutations in cancer for both proteins, including the highlighted arginines in LDOC1, which form electrostatic interactions with the opposite chain. TWIST1 has several annotated pathogenic mutations, including L149R and L159H, which are at residues buried in the interface (**Fig. 4F**). In particular, the leucine to arginine mutation, associated with the Saethre-Chotzen syndrome, would strongly disrupt packing. The R118G mutation would disrupt the interaction with residue F22 mainchain O in TCF4. In RAD51D we found the mutation R266C (Breast-ovarian cancer, familial) that interacts across the interface with XRCC2 (**Fig. 4G**), paralogous genes involved in the repair of DNA double-strand breaks by homologous recombination. Interestingly, we also found mutations at R239, to Trp/Gln/Gly, associated with Breast-ovarian cancer that interacts with Tyr119 in XRCC2 that itself is also annotated as having mutations linked to hereditary cancer-predisposing syndrome.

### Phospho-regulation of protein complex interfaces

Protein phosphorylation can regulate protein interactions by modulating the binding affinity via the change in size and charge of the modified residue. Over 100,000 experimentally human phosphorylation sites have been determined to date ^26,27^, but only 5 to 10% of these have a known function ^28^. Mapping phosphorylation site positions to models of protein interfaces can generate mechanistic hypotheses for the functional role of phosphorylation sites in controlling protein interactions. We used a recent characterisation of the human phosphoproteome ^26^ to identify 4145 unique phosphosites at interface residues of the set of confidently predicted structures. We noted that the average functional importance, defined by the functional score described by Ochoa and colleagues ^26^, was generally higher than random for phosphorylation sites at interfaces (**Fig. 5A**). Among the interface phosphorylation sites, we found some enrichment for targets of multiple kinases, including several tyrosine kinases (ERBB2, AXL, ABL2, FER) (**Fig. 5B**). This observation suggests that some interfaces in different protein pairs may be under coordinated regulation by specific kinases and conditions.

**Figure 5.**
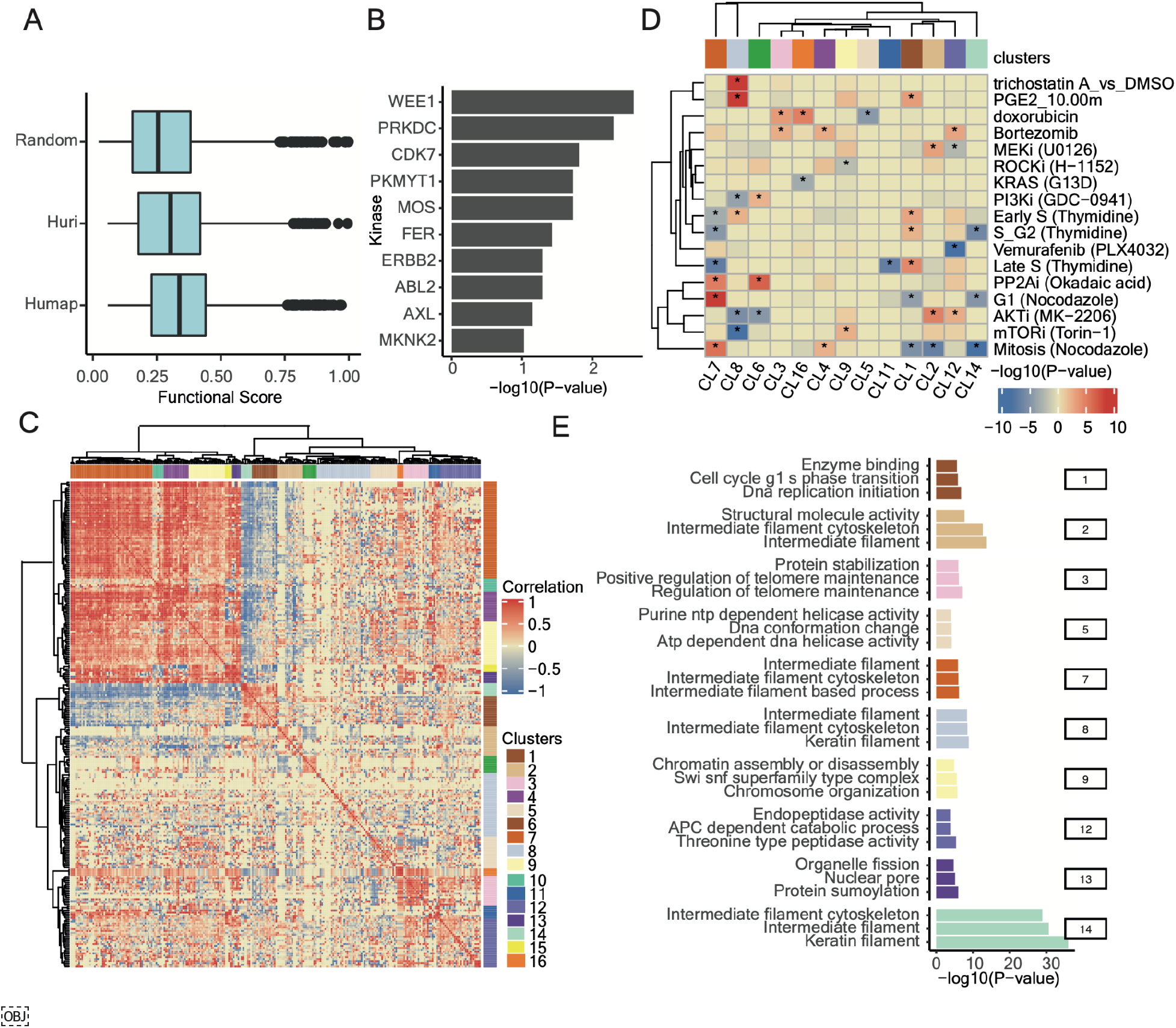
Co-regulation of phosphorylation sites at interface residues. A) Distribution of phosphosite functional scores for phosphosites at interface residues and random phosphosites. Enrichment of kinase substrates among phosphosites at interface residues. C) Hierarchical clustering of the pairwise correlation values for changes in phosphosites levels across conditions. Groups of phosphosites showing high correlation values were defined as clusters (1 to 16) as indicated in colours along the outside of the clustergram. D) Degree of regulation of phosphosites from each cluster in a select panel of conditions, defined as a Z-test comparing the fold change of the phosphosites in a cluster compared with the entire distribution of fold changes in that condition. The result is summarized as the -log(P-value) and signed as positive if the median value is above the background or negative otherwise. E) Gene-ontology enrichment analysis for the proteins with phosphosites annotated to select clusters.

To identify potentially co-regulated interfaces, we collected measurements of changes in phosphorylation levels across a large panel of over 200 conditions ^29^. We retained 260 phosphosites that had a significant regulation in three conditions and then computed all-by-all pairwise correlations in phosphosite fold changes across conditions. We clustered these phosphosites by their profile of correlations (**Fig. 5C**), identifying 16 groups of co-regulated interface phosphorylation sites (**Fig. 5C, Table S3**). For each group of phosphosites, we identified the conditions where these have the strongest up- or down-regulation (**Fig S1**) and plotted a subset of conditions in **Fig. 5D**. We also performed a GO enrichment analysis for each group of co-regulated phosphosites, including both proteins of the modified interfaces, to search for common biological functions (**Fig. 5E, Table S4**). For example, we observed a cluster of interface phosphosites in proteins related to intermediate filaments (cluster 7) that show strong regulation patterns along the cell cycle, downregulated in S-phase and up-regulated in G1 and mitosis. Phosphosites in cluster 1 (cell cycle G1-S phase transition) show the opposite trends with up-regulation in late S-phase and down-regulation in G1 and mitosis. Some clusters show regulation under specific kinase inhibition which may provide novel hypotheses for kinase regulation of specific processes. For example, phosphosites in cluster 9 (regulation of chromosome assembly) tend to be up-regulated after inhibition of ROCK and up-regulation after inhibition of mTOR.

While not all phosphosites at interfaces are likely to regulate the binding affinity, this analysis provides hypotheses for the potentially coordinated regulation of multiple proteins by tuning their interactions after specific perturbations. We provide the complete list of interface phosphosites, known kinase regulators and condition-specific regulation in **Table S1**.

### Higher-order assemblies of protein complexes from binary interactions

Proteins interact with multiple partners either simultaneously, as part of larger protein complexes, or separated in time and space. This is also reflected in our structurally characterised network, where proteins can be found in groups as illustrated in a global network view of the interactions with confident models (**Fig 6**, central network, **Fig S2 and Data S1**). One key benefit of structurally characterising an interaction network is the identification of shared interfaces for multiple interactors. As an example, we highlight GDI1 (Rab GDP dissociation inhibitor alpha) that interacts with multiple Rab proteins regulating their activity by inhibiting the dissociation of GDP. The predicted complex structures for these interactions shows how these share the same interface and therefore cannot co-occur. Other clusters in the network suggest that the proteins form larger protein complex assemblies with many-to-many interactions. As the use of AlphaFold2 for predicting larger complex assemblies can be limited by computational requirements, we tested whether the structures for pairs of proteins could be iteratively structurally aligned. We tested this procedure on a small set of complexes covered in this network, with known structures and the number of subunits ranging from 5 (RFC complex, TFIIH core complex) to 14 (20S proteasome). We then aligned an experimentally determined structure with the predicted models (**Fig. 6**, grey - experimental model). These examples showcase the potential and also limitations of this procedure.

**Figure 6.**
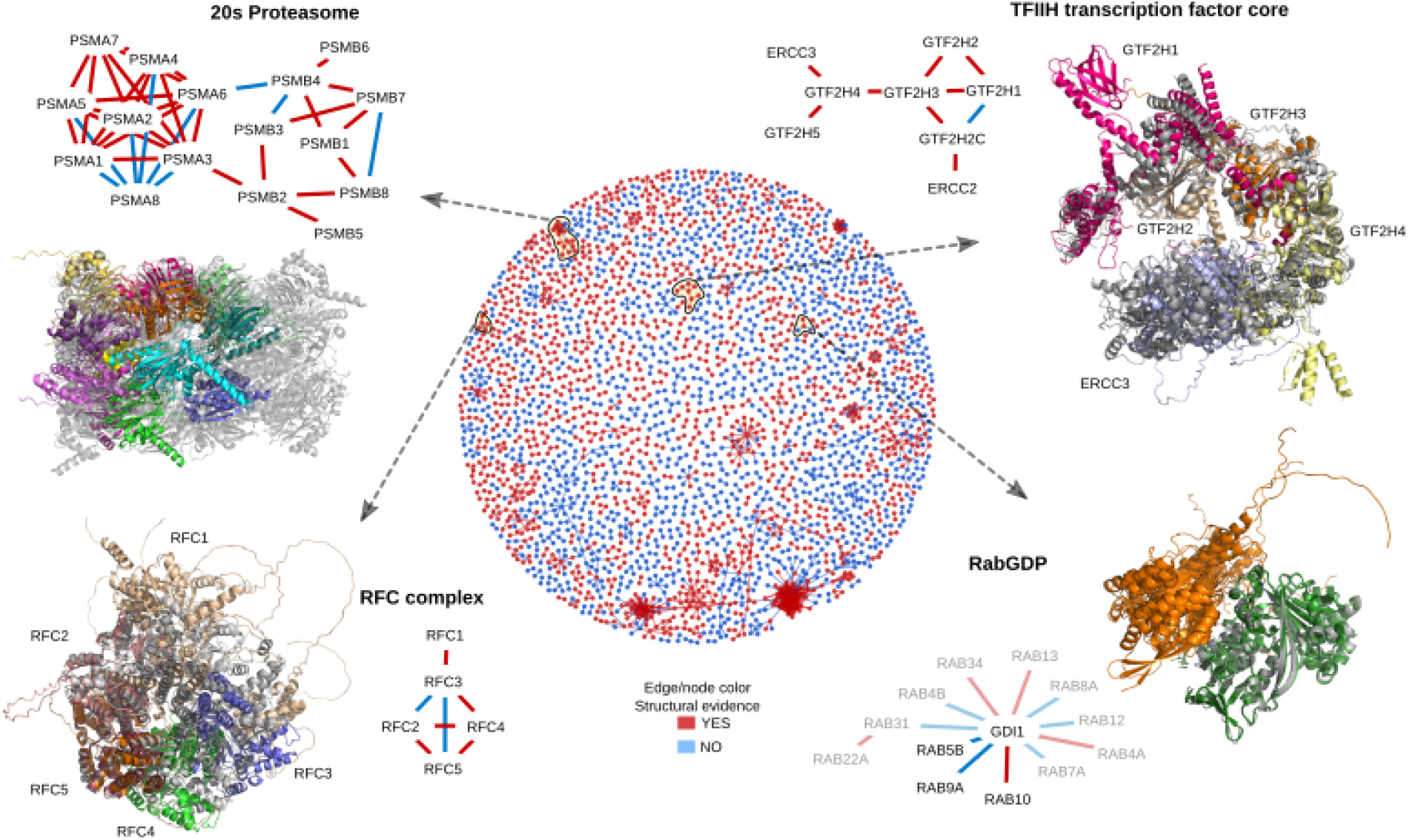
Protein complex predictions for higher-order assemblies. The middle circle is a network view of all protein-protein interactions predicted with high confidence (pDockQ>0.5). The edges and nodes are coloured in red if there is a previous experimental or homology model for the interaction, in blue if such information is not available. We selected four examples of recapitulated complexes (yellow circles and black arrows) plotted in further detail. In these small networks, only the edges are coloured based on structural evidence. In the case of RabGDP the faded nodes and edges represent predictions with slightly lower confidence (pDockQ>0.3)

The TFIIH core complex is composed of 5 subunits with 1-to-1 stoichiometry. All subunits can be modelled with the final complex generally agreeing (**Fig. 6**) with a cryoEM structure for these subunits (6NMI). The most significant difference to the cryoEM model is the relative positioning of the ERCC3 subunit. The exact final model obtained can vary depending on the aligned pairs with multiple possible final conformations (**Fig S3**). **Fig. 6** illustrates the conformation that best matches the cryoEM model in 6NMI. For example, for the TFIIH core complex, there is a predicted model where the complex adopts a more open conformation (as seen in 5OQJ) and alternative predicted placements of the GTF2H1 subunit.

The RFC complex is also composed of 5 subunits with 1-to-1 stoichiometry. One iterative alignment of pairwise interactions builds a model that includes all five subunits organised similarly to that observed in the 6VVO cryoEM structure (**Fig. 6**). In this predicted model, the subunits RFC2/5/4/3 match the experimentally observed model well, but there are apparent deviations introduced by compounding errors in alignment by this iterative process. Individual subunits in the cryoEM structure can be aligned to each of the model subunits well, but then the alignment of the rest of the model is progressively worse the further away the subunits are positioned from the aligned subunit. The RFC1 subunit is individually not well predicted, showing a considerable difference between the cryoEM and AlphaFold2 models. Some of the modelled pairs highlighted additional issues. For example, the RFC3 - RFC5 interaction pair is predicted with high confidence, while in fact, these do not share a direct contact in the experimental structure. AlphaFold2 places RFC3 at the RFC5-RFC4 interface, likely due to the structural similarity between RFC3 and RFC4.

Encouraged by the examples tested, we defined an automatic procedure to generate larger models by iterative alignment of pairs (**Methods**). We start building all possible dimers in a complex, then sort them by pDockQ, and start building from the first ranked dimers. Next, we add the highest-ranked dimer, which shares one subunit with the complex if it does not overlap; this is repeated for all dimers until the complex is complete or no additional proteins can be added. We tested this on the 20S proteasome, a particularly challenging example with stoichiometries different from 1-to-1 and homologous subunits. This automatic procedure could build a model containing all 14 subunits (half of the proteasome) that are mostly placed in agreement within the experimental model (**Fig. 6**). However, the exact order of the chains is incorrect, i.e. at each location an incorrect protein is placed, highlighting that AF2 cannot distinguish which two proteins interact from a set of homologous proteins.

Two additional proteins where we could build a good model are Heterodisulfide reductase from Methanothermococcus thermolithotrophicus (pdb:5ODC) and the eukaryotic translation initiation factor 2B from Schizosaccharomyces pombe (5B04), see supplementary **Fig S4**. For 5ODC we could build a complete model of the protein with an RMSD of 6.0Å (TM-score 0.90) starting from dimers. However, for 5B04 it was not possible as the chains started overlapping when we tried to build a larger model. However, if we build trimers and then use all three dimers from these trimers we can build a complete model with an RMSD of 7.3 Å (TM-score 0.86), showing that it is sometimes necessary to use largers subunits to assembly the complexes. Preliminary results from a follow up study show that it is often possible to build the structure of several complexes if the subunits are well predicted. We have developed a computationally efficient tool for this method using a Monte Carlo Tree search^30^. In summary, we find that it is possible to iteratively align structures of pairs of interacting proteins to build larger assemblies but identified issues that limit this procedure at the moment.

## Discussion

We have attempted here to generate predicted complex structures for pairs of human proteins known to physically interact from two different datasets based on different experimental approaches. We noted that the source of data used for the protein interactions is important and impacts the fraction of models that can be confidently predicted. Our analysis suggests the protein interactions supported by a combination of affinity, co-fraction, and complementation based methods results in higher confidence models. We believe these interactions tend to correspond to high-affinity interactions that are very likely to share a direct physical permanent interaction. We show that it is possible to use metrics from the models (e.g. pDockQ score) to rank higher confidence models, providing an additional accuracy level to the large scale protein-protein interaction studies. Further, future large-scale computational predictions of protein-protein interactions may provide additional high-quality targets for detailed studies of stable complexes. Experimental data from cross-link mass spectrometry experiments provide an ideal resource for further validating these predictions via orthogonal means. In principle, such constraints from cross-link could also be considered during predictions, and it may be possible in the future to develop predictors that can take in such constraints as part of the starting information.

Based on comparisons with solved structures, we suggest that models with pDockQ>0.5 are very likely to be correct. Additionally, models with lower scores (0.5>pDockQ>0.23) are still likely to contain many correct solutions and may highlight correct interfaces even if not fully correct orientations of the interacting proteins, in agreement with results from CAPRI^31,32^. In this study this would correspond to an additional 6000 complex structures. Such lower confidence models are likely to be useful for generating hypotheses and large-scale analysis of global properties. Equally important is the caveat that high confidence predictions will still contain errors, and in particular, we note that in protein complexes containing paralogous proteins (which is common in higher eukaryotes^33^), the current procedure cannot identify the exact pairing of the protein. For such cases, additional methods need to be developed.

Structural models for protein interfaces are critical for understanding molecular mechanisms and the impact of mutations and post-translational modifications. We illustrate this using disease mutations and phosphorylation data. While much disease-associated variation is often found in non-coding regions of the genome, the growth of exome sequencing of large cohorts of patients will lead to discovering many more protein mutations linked to disease, which will require such large structural characteristics. Both for mutations and phosphorylation sites, we think these analyses should be seen as generating hypotheses for further testing, and we make this information available in the supplementary material to facilitate such future work.

Finally, in principle, we show that it is possible to build structural models for larger assemblies from the binary complexes predicted here. In a follow-up paper we have shown that it is sometimes possible to build large assemblies fully automatically by using predictions of dimers and trimers^30^. Aspects that may limit this include the structural homology between subunits, unknown subunit stoichiometries and limits in the predicted interactions^30^. Additional work will be needed to determine the exact stoichiometry and design methods and score systems to build such larger complex assemblies, as well as to predict the interaction of proteins with weak and transient interactions.

## Methods

### Protein interaction data and annotations

Human protein pairs known to physically interact were obtained from the Hu.MAP dataset, retaining pairwise interactions with >=0.5 confidence, and most interactions from the HuRI dataset. These interactions were further enriched by obtaining annotations on cross-linked peptides matched across pairs of interaction proteins, disease related mutations and protein phosphorylation sites in the selected proteins. In addition all non homologous pairs from 12 protein complexes (Table S5) and 4320 protein pairs from 2102 different protein complexes (Table S6) in CORUM^23^ were used for additional analysis. A complete list of all datasets is available from the supplementary data. A subset of Cross-link data was collected from ^34–44^, filtered for peptides unique to only one protein sequence. A cross link was considered validated by the structure if the distance between the epsilon amino groups on the side-chains of the relevant pair of lysine residues were within 32Å. Clinical missense variants associated with disease were collected from ClinVar. We selected only those having pathogenic or likely pathogenic effects which were mapped to Uniprot protein sequences using VarMap. The final list of mutated positions was then compared to the interface positions. We obtained a list of protein phosphorylation sites with predicted functional relevance ^26^, phosphosite annotations ^28^ and regulation of phosphorylation sites across a large panel of conditions ^29^. These phosphosites were also mapped to interface positions as defined by the predicted models. All protein interaction networks were processed using R packages igraph (v1.2.5) and qgraph (v 1.9), further graphical editing was done using Cytoscape ^45^.

### Protein complex prediction

To predict protein complexes of pairwise interactions, we used the FoldDock pipeline ^18^ based on AlphaFold2 ^17^. We use the option of fused+paired multiple sequence alignments (MSAs) and run the model configuration m1-10-1 as this provides the highest success rate accompanied by a 20-fold speed-up. Both the fused and paired MSAs are constructed from running HHblits on every single chain against Uniclust30. The fused MSA is generated by simply concatenating the output of each of the single-chain HHblits runs for two interacting chains. The paired MSA is constructed by combining the top hit for each matching OX identifier between two interacting chains, using the output from the single-chain HHblits runs.

### pDockQ confidence score

To score models, we use features from the predicted complexes to calculate the predicted DockQ score, pDockQ. This score is defined with the following sigmoidal equation:

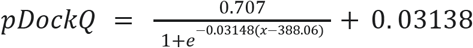

where,

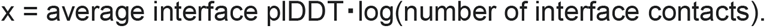

The parameters were optimised to predict the DockQ score using the dataset from ^46^. The number of interface contacts is defined as elsewhere in this paper (any residues with an interface atom within 10Å to the other chain), and the plDDT is the predicted lDDT score from AlphaFold2 taken over the interface residues as defined by the interface contacts.

### Building larger complexes from binary interactions

A simple procedure to build larger complexes from a set of paired models was developed. All dimers in the set are by default ranked by their pDockQ values.

1. The building is started from a single dimer by default the dimer with the highest pDockQ value. This is referred to as the “complex”.
2. All other dimers in the set are then tried to be added to the “complex. Starting with the one with the second highest pDockQ a chain is added to the complex if:
  a. Exactly one chain of the dimer is identical to one chain in the complex
  b. The structure of these two chains is similar enough (default TM-score > 0.8)
  c. The dimer is then rotated so that the two chains overlap-
  d. The second chain in the dimer does not clash with more than 25% of its residues (CA<5Å) to any chain in the complex.
3. If a chain is added, the procedure is started over again and repeated until no more chains can be added.

### Analysis of phosphosites in the protein-protein interfaces

Phosphosite residues in interfaces were identified from a previously published comprehensive list of known human phosphosites ^26^. Kinases associated with phosphorylation of interface residues were obtained from the PhosphositePlus database and over-representation analysis of kinases was performed using a hyper-geometric test. Highly regulated interface phosphosites were defined as those with more than two-fold change in phosphorylation in more than two perturbation conditions across a collated phosphoproteomics dataset comprising a range of physiological conditions and drug treatments ^29^. Pearson correlation was calculated amongst these regulated phosphosites and clusters of co-regulated phosphosites were identified using hierarchical clustering (‘ward’ method) of Euclidean distances of the correlation matrix. Phosphosite clusters were created by cutting the dendrogram at the appropriate level using the cutree (h=17) function in R. Phosphosite clusters that were significantly regulated in each perturbation condition were identified by a z-test from the comparison of fold changes in phosphosite measurements of all phosphosites in a cluster against the overall distribution of phosphorylation fold changes across the condition. Gene ontology over-representation of each cluster was performed separately using a hypergeometric test in R. The gene ontology terms were obtained from the c5 category of Molecular Signature Database (MSigDBv7.1) ^47^. All over-representation analysis (ORA) were performed using the enricher function of clusterProfiler package (version 3.12.0) ^6^ in R.

## Comparison with other databases

All proteins used here were mapped to UniProt^48^ to retrieve subcellular localisation, STRING^49^ for coexpression and other interaction data, gtex^50^ for tissue specific expression.

## Availability

All code used in this project can be found at https://gitlab.com/ElofssonLab/huintaf2/. Tools to run AlphaFold2 can be found at https://gitlab.com/ElofssonLab/FoldDock/. All models generated as well as some of the multiple sequence alignments can be found at https://archive.bioinfo.se/huintaf2/. All datasets and meta-data is available from

## Acknowledgements

R.A.S. acknowledges funding through the European Union Horizon 2020 program INFRAIA project Epic-XS (Project 823839) and the research programme NWO TA with project number 741.018.201, which is partly financed by the Dutch Research Council (NWO). AE was funded by the Vetenskapsrådet Grant No. 2016-03798 and Knut and Alice Wallenberg foundation. The computations/data handling were enabled by the supercomputing resource Berzelius provided by National Supercomputer Centre at Linköping University and the Knut and Alice Wallenberg foundation and SNIC, grant No: SNIC 2021/5-297 and Berzelius-2021-29. J.E.B. acknowledges funding from the National Heart Lung and Blood Institute (NHLBI 5R35GM13625) and National Institute for General Medical Sciences (NIGMS 5R01HL144778).

