## Supplementary figures and images for "Towards a structurally resolved human protein interaction network"

### Figure S1

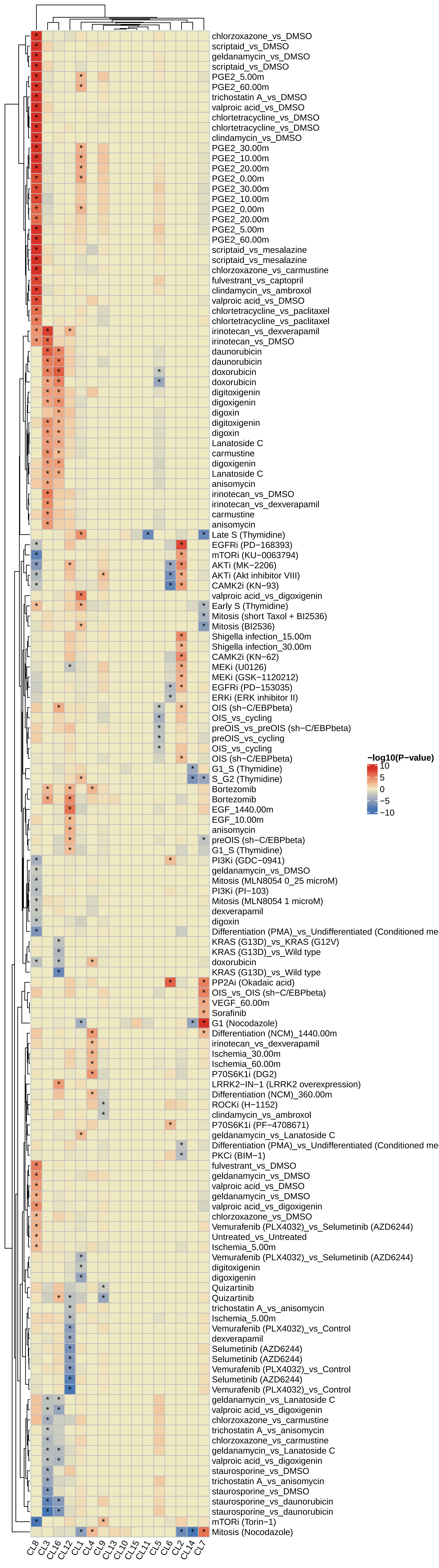

### Figure S2

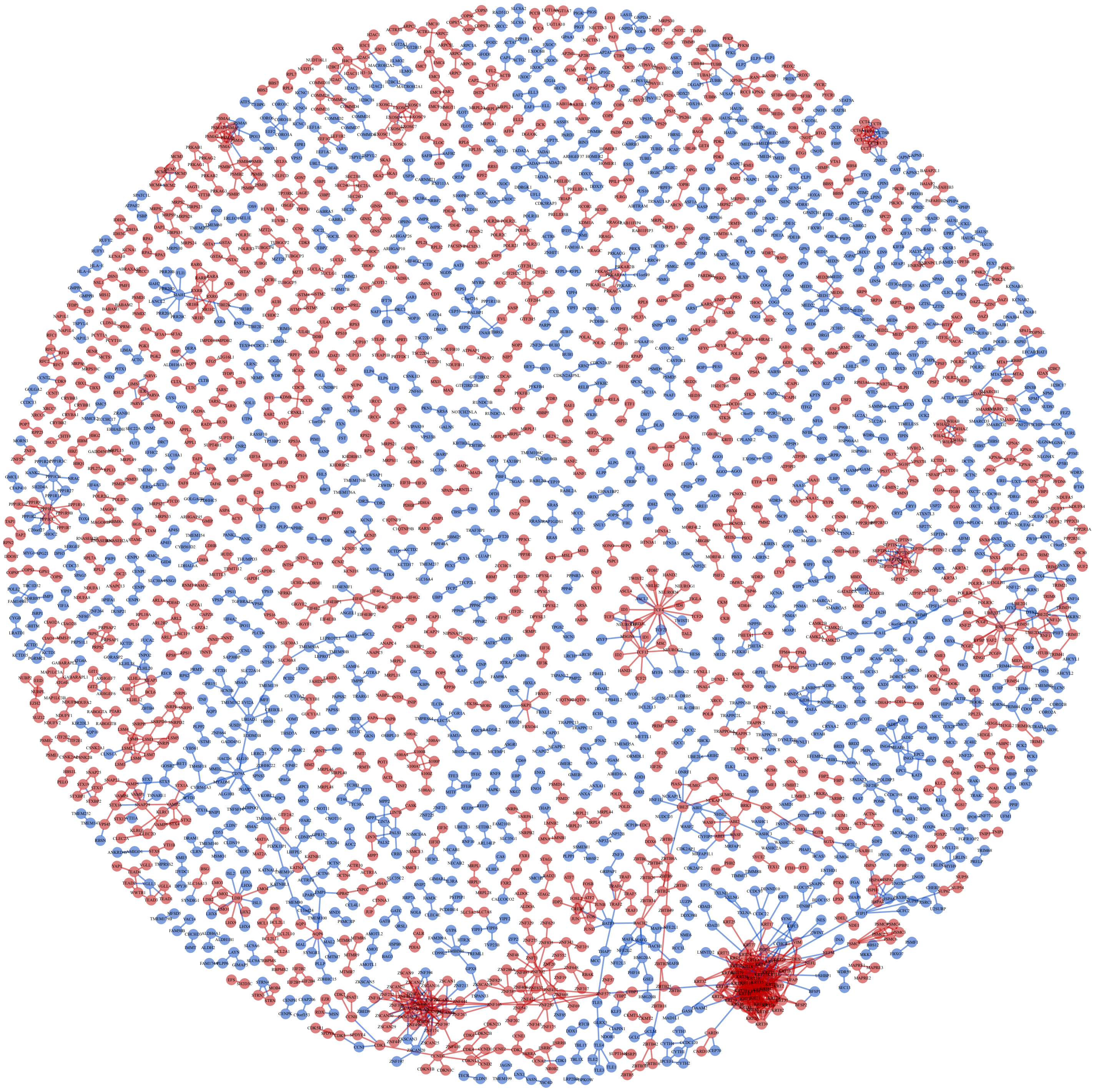

### Figure S3

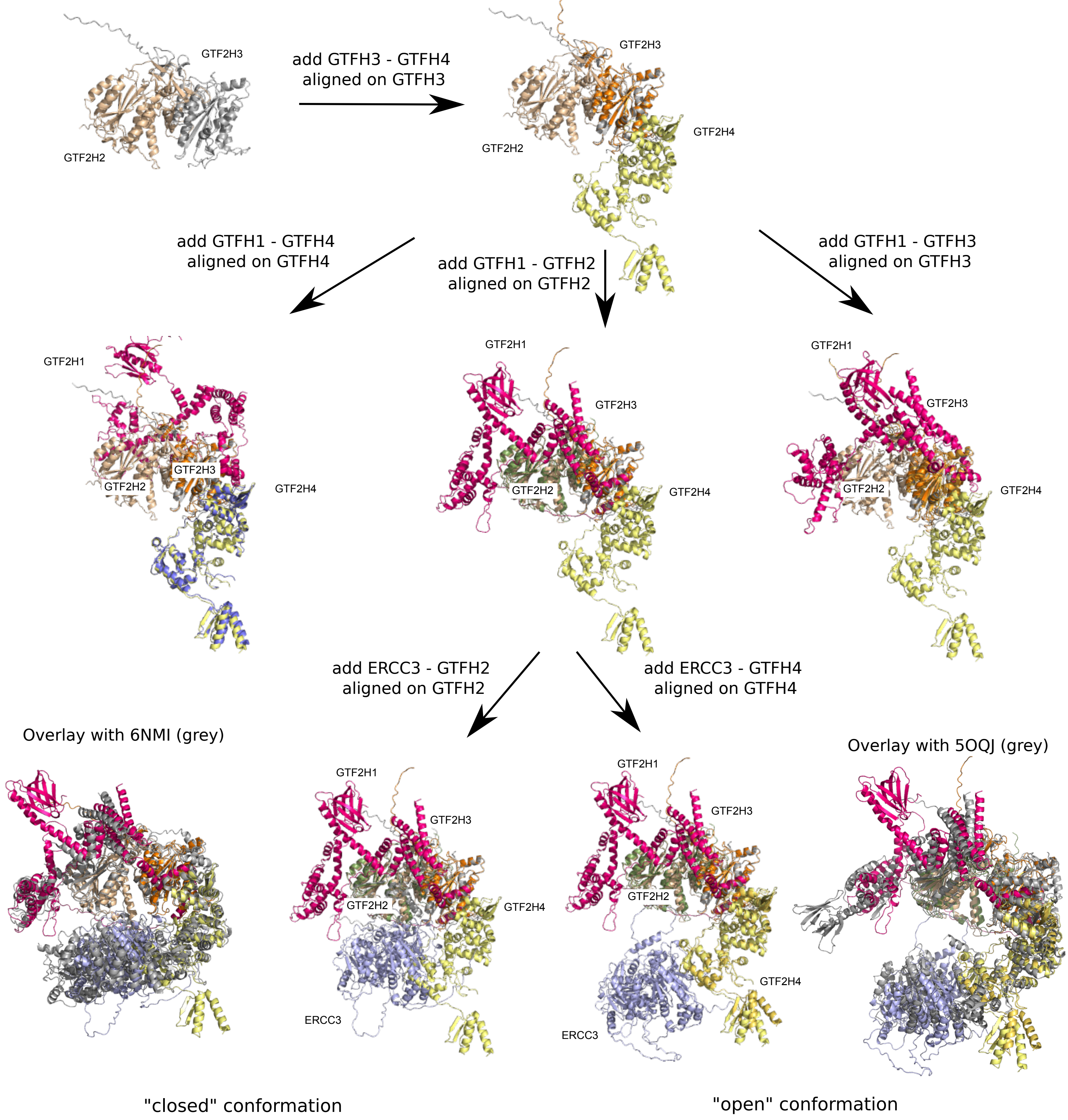

### Figure S4

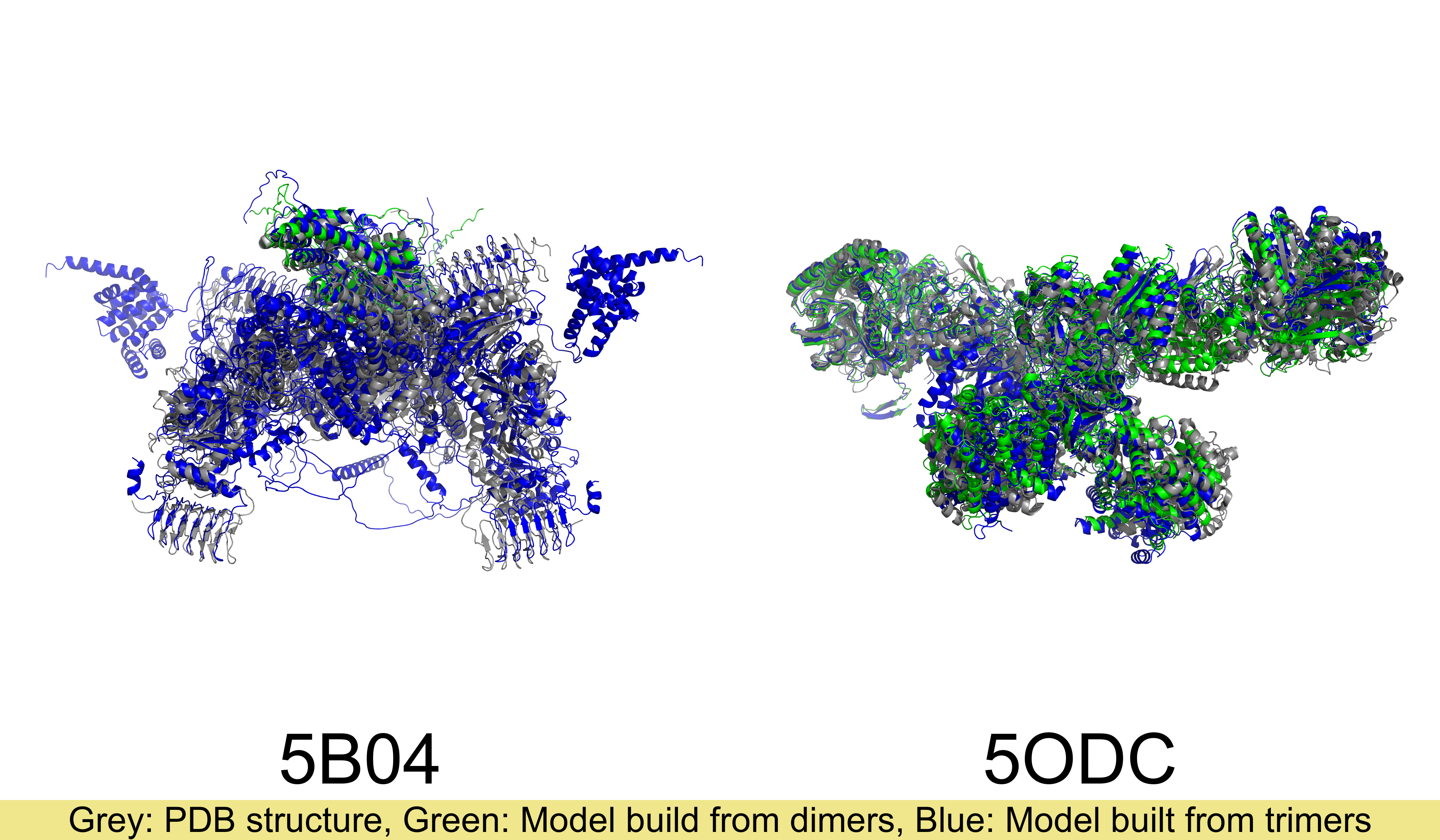
